# A study on packed cell volume for deducing hemoglobin: Cholistani camels in perspective

**DOI:** 10.1101/2022.11.15.516630

**Authors:** Umer Farooq, Musadiq Idris, Mushtaq Hussain Lashari, Shahbaz Ahmad, Nouman Sajjad, Zia-Ur-Rehman, Haroon Rashid, Aisha Mahmood, Sajid Hameed

**Affiliations:** Department of Physiology, The Islamia University of Bahawalpur, Pakistan; Department of Zoology, The Islamia University of Bahawalpur, Pakistan; Department of Anatomy and Histology, The Islamia University of Bahawalpur, Pakistan

**Keywords:** packed cell volume, hemoglobin, Cholistani camels

## Abstract

In human medical practice, a hematological rule of three has been validated for healthy human populations. One such formula is estimating hemoglobin (Hb) levels as 1/3^rd^ of Packed Cell Volume (PCV). However, no such hematological formulae have been devised and validated for veterinary medical practice. The present study was devised with an aim to evaluate the relationship between hemoglobin (Hb) concentration and Packed Cell Volume (PCV) in camels (n=215) being reared under pastoralism, and to devise a simple pen-side hematological formula for estimation of Hb from PCV. The PCV was determined through microhematocrit method whereas Hb estimation by cyanmethaemoglobin method (HbD). The Hb was also calculated as 1/3^rd^ of PCV and was dubbed as calculated Hb (HbC). Overall HbD and HbC were significantly (P≥0.05) different. Similar results were attained for all study groups *i.e*. males (n=94) and females (n=121), and young (n=85) and adult (n=130) camels. The corrected Hb (CHb) was deduced through regression prediction equation attained from linear regression model. Scatter-plots were drawn, linear regression was carried out, and Bland Altman chart was built for agreement of both methods of Hb estimation. A non-significant (P≥0.05) difference was noticed between HbD and CHb. Bland Altman chart revealed good agreement between HbD and CHb and there was no proportional bias on the distribution of data around the mean difference line (Mean= 0.1436, 95% CI= 3.00, -2.72). We accordingly recommend a simplified pen-side hematological formula for deducing Hb concentration from PCV *viz*. Hb concentration (g/dL)=0.18(PCV)+5.4 for all age and gender groups of camels instead of its calculation as one-third of PCV.

## Introduction

Packed Cell Volume (PCV), also well-known as hematocrit or erythrocyte volume fraction, is fraction of red blood cells (RBCs) in the animal’s blood [1]. PCV is responsible for transportation of oxygen and absorbed nutrients [2]. Amplified PCV not only results in a better transportation but also an augmented primary and secondary polycythemia [3]. Moreover, a high PCV reading pointed out either an increased number of RBCs or decreased volume of circulating plasma. PCV is the most precise way of estimating erythrocyte volume and may also be used to assume total blood volume and hemoglobin (Hb) level.

The manual, spun PCV (through microhematocrit method) is a key measurement, underpinning much of hematology. The calibration of virtually all hematology autoanalyzers can be traced in some way back to the PCV [4]. Reference ranges for the hematocrit and red cell indices depend on the validity of this calibration, as do the assignment of expected values to calibrators and controls, and the assignment of target values for statistical population-based quality control programs. Any errors in PCV assignment have far-reaching implications [5].

Extensive research work has been conducted in human medical sciences directed towards assessing an interrelationship between PCV and Hb, and confirming the thumb rule of Hb being 1/3^rd^ of PCV. Certain studies have nullified this rule claiming that Hb estimates cannot be obtained from PCV values with a reliable precision by making use of the common rule of dividing by three [6, 7, 8]. The results of these studies also indicated that the association between Hb and PCV is not exactly three times and the sex and age of the individuals can also have a significant effect on this three-fold conversion.

For veterinary medical sciences, the interrelationship between PCV and Hb has been studied for indigenous cattle [9]. In this study assessing Hb as 1/3^rd^ of PCV has been nullified and a newer alternative formula has been reported for Hb assessment through PCV. Similarly, our laboratory has reported similar reports and pen-side hematological formulae for Hb estimation for goats [10] and Cholistani breed of cattle [11]. However, study on such interrelationship for the blood of camels has not yet been reported. The present novel study has therefor been devised with an aim of evaluating the relationship between Hb concentration and PCV in Cholistani camels being reared under pastoralism in Cholistan desert of Pakistan. Furthermore, it also aims to devise a simple pen-side hematological formula for estimation of Hb from PCV.

## Materials and Methods

### Geo-location of the Study

The study was simultaneously conducted at Cholistan desert, Pakistan (for field blood sampling) and Physiology post-graduate lab of the Department of Physiology, The Islamia University of Bahawalpur (IUB), Pakistan (for lab work). Cholistan desert is located at latitudes 27°42’and 29°45’North and longitudes 69°52’and 75°24’East and at an altitude of 112m above the sea level. The climate of this area is arid, hot subtropical and monsoonal with the average annual rainfall of 180 mm. The mean annual temperature is 28.33°C, with the month of June being the hottest when the daily maximum temperature normally exceeds 45°C [12, 13].

### Experimental Animals

Cholistani camels (n=215) were randomly selected from nomadic pastoralists for blood sampling irrespective of their age and sex. All the animals were being reared under similar management and feeding conditions of pastoralism either under transhumanie or nomadic pastoral livestock production systems [14]. Split-herding is normally exercised for livestock by the pastoralists, according to which the young ones (calves in this case) are kept at their pens near the ‘Tobas’ (natural or man-mad water reservoirs of the desert), while the adults are sent for grazing till night-time [12]. The general health status of animals was ascertained through a thorough anamnesis from the livestock owners and clinical signs. The animals which were found to be lethargic, depressed, off-feed and segregated from the herd (as per the anamnesis taken from the pastoralist herders) were not included in the study.

### Blood Collection

The blood sampling was conducted from July to October, 2022. About 5 mL blood was collected aseptically in anticoagulant-added tubes (0.5 M EDTA) with the help of a disposable syringe from the high neck jugular vein of each animal. The same restraining technique with same personnel and time were used to minimize the stress in animal and also to normalize blood collection procedure. The blood samples were mixed by gentle inversion and transported in an ice box to the Physiology Post-graduate Lab, IUB, Pakistan, refrigerated and analyzed within 24h for hematological analyses.

### Hematological Analyses

The blood samples were analyzed for PCV and Hb as per the protocols prescribed by the WHO and in vogue, and are considered as gold standard tests for PCV and Hb determination, respectively [1]. PCV was deduced by microhematocrit centrifuge method using microcentrifuge (Sigma Aldrich, Model 5254, Germany) and reading as percentage (%) was taken through a hematocrit card-reader. The reading was used for calculating Hb as its third and was dubbed as Hemoglobin Calculated (HbC).

The Hb concentration was also determined through Drabkin’s reagent (HbD) using cyanmethaemoglobin method and a commercial Hb Kit (AMP Diagnostics, BD6100-E V4.0-CE Ameda Labordiagnostik GmbH, Germany) [15]. The Hb was calculated as per formula prescribed in instructions manual.

### Statistical Analyses

Statistical Package for Social Science (SPSS for Windows version 12, SPSS Inc., Chicago, IL, USA) was used for data analysis. Means (±SE) and 95% CI for hematological attributes (PCV and Hb) were computed using prescribed formulae. For the purpose of analyses, considering the non-normal nature of the attained data, the Mann Whitney-U test was implied as a non-parametric test for deducing difference between HbD and HbC, and between HbD and corrected Hb (CHb) for all study groups (young= 85, adult= 130; females= 121, males= 94).

Linear regression analyses were carried out, scatter-plots were drawn between the following as prescribed earlier and Bland Altman test was implied [9, 16]:

a. HbD and PCV
b. HbD and HbC
c. The difference of HbD and CHb (HbD-CHb), and means of measurements (HbD+CHb/2)

Regression prediction equations were accordingly computed. Attaining a highest adjusted r-square value from these equations, CHb was calculated.

## Results

In the present study, hemoglobin determined spectrophotometrically (HbD) and hemoglobin calculated (HbC) as one third of Packed Cell Volume (PCV) was assessed for statistical difference at P≤0.05. Furthermore, PCV conducted through microhematocrit method was studied for interrelationship between the HbD and corrected hemoglobin (HbC) (through a formula attained by regression analyses).

Regarding normality of studied attributes (HbD, PCV and HbC), the Shapiro-Wilk test revealed that all the three studied attributes were not distributed normally.

Mean (±SE) values and 95% CI for hematological attributes (HbD, HbC and PCV) in Cholistani camels (n=215) are presented in Table 1. The overall results indicated a significant (P≤0.05) difference between HbD and HbC. Furthermore, similar results were attained for all study groups (females *vs* males, and adults *vs* young) of the present study.

**Table 1.** Mean (±SE) values and confidence intervals for hemoglobin determined (HbD), hemoglobin calculated (HbC) and Packed Cell Volume in Cholistani camels (n=215)

| Groups |  | HbD |  | HbC |  | Sig | PCV |  |
| --- | --- | --- | --- | --- | --- | --- | --- | --- |
|  |  | (g/dL) |  | (g/dL) |  |  | (%) |  |
|  |  | x±SE | CI | x±SE | CI |  | x±SE | CI |
| Gender | Females (n=121) | 10.8±0.2 | 10.4-11.2 | 9.4±0.2 | 9.0-9.8 | 0.01* | 28.3±0.6 | 27.2-29.5 |
|  | Males (n=94) | 11.2±0.3 | 10.7-11.8 | 10.8±0.4 | 10.0-11.7 | 0.04* | 32.6±1.2 | 30.1-35.1 |
| Age | Young (n=85) | 11.0±0.3 | 10.3-11.6 | 9.3±0.5 | 8.2-10.2 | 0.005* | 27.8±1.4 | 24.8-30.7 |
|  | Adult (n=130) | 11.1±0.2 | 10.7-11.5 | 10.4±0.3 | 9.8-11.0 | 0.05* | 31.3±0.8 | 29.6-32.9 |
| Overall (n=215) |  | 11.0±0.1 | 10.7-11.4 | 10.2±0.3 | 9.7-10.7 | 0.007* | 30.7±0.7 | 29.2-32.1 |

\*Significant at $P \leq 0.05$ within rows for each group between hemoglobin determined and hemoglobin calculated.
HbD= Hemoglobin determined spectrophotometrically; HbC= Hemoglobin calculated as $1/3^{\text{rd}}$ of PCV; PCV= Packed cell volume

The results for linear regression for all study groups are presented in Table 2. Significantly (P≤0.01) higher positive correlation coefficient was noticed for young camels (r=0.830; adjusted r-square=0.68) between HbD and PCV, and between HbD and HbC.

Regression equations were developed to validate the 1/3^rd^ association between PCV and HbD for all study groups. The overall results regression equation hence attained *i.e*. Hb (g/d)= 0.18(PCV)+5.4 was used to deduce Hb dubbed as corrected Hb (CHb). A non-significant (P≥0.05) difference was noticed between HbD and CHb. This equation is therefore, considered valid for deducing Hb from PCV in all age and gender groups of Cholistani camels.

The scatterplots of spectrophotometrically determined Hb (HbD) versus PCV, and HbD versus hemoglobin calculated as 1/3^rd^ of PCV (HbC) have been given in Figure 1 (a,b). Similarly, the scatterplots and Bland and Altman Chart for difference between HbD and CHb (HbD-CHb) versus average of HbD and CHb (HbD+CHb/2) is given in **Error! Reference source not found.**2. Bland Altman chart revealed good agreement between HbD and CHb and there was no proportional bias on the distribution of data around the mean difference line (Mean= 0.1436, 95%CI= 3.00—2.72).

**Table 2.**
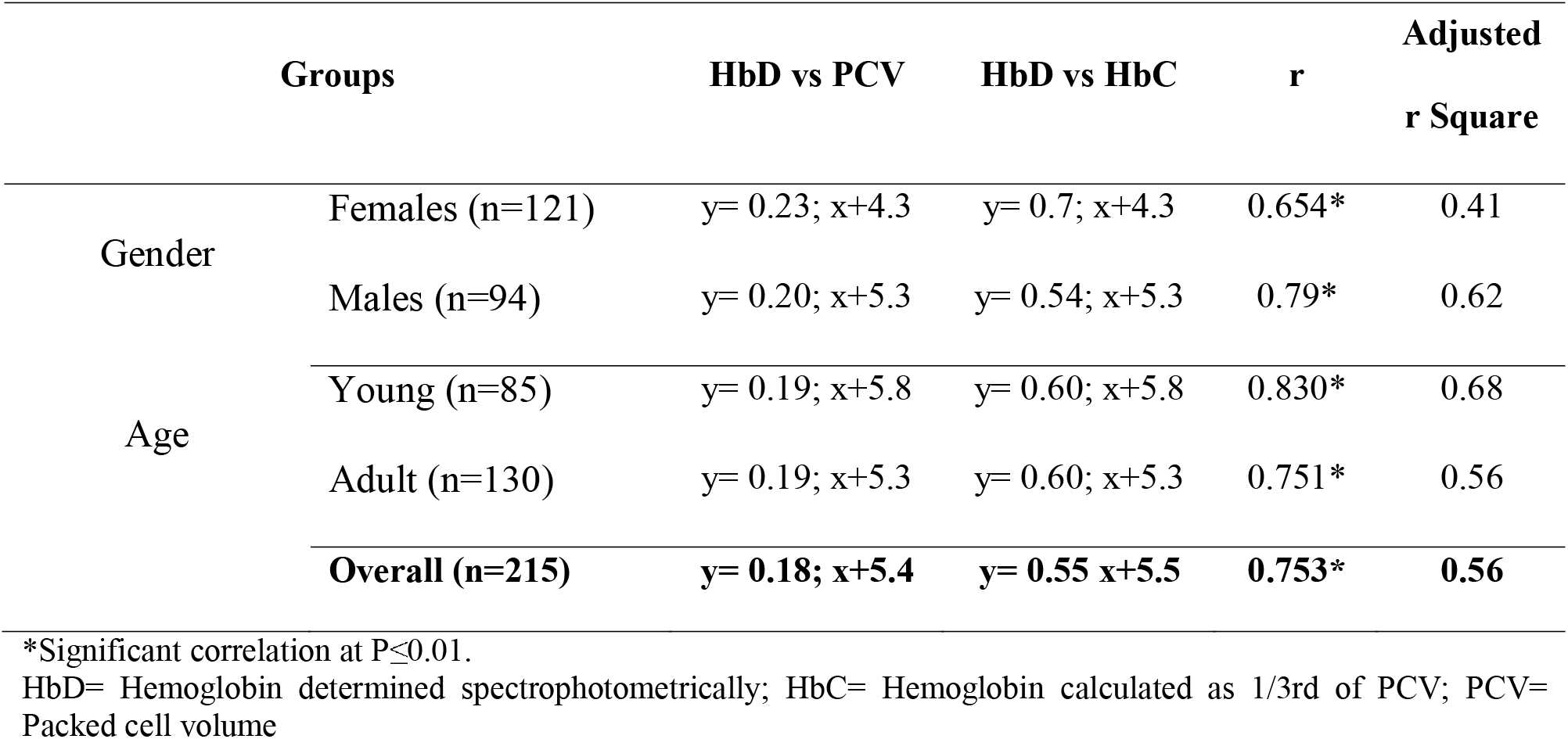
Linear regression between various hematological attributes for Cholistani cattle (n=215)

| **Groups** |  | **HbD vs PCV** | **HbD vs HbC** | **r** | **Adjusted**<br>**r Square** |
| --- | --- | --- | --- | --- | --- |
| **Gender** | Females (n=121) | y= 0.23; x+4.3 | y= 0.7; x+4.3 | 0.654* | 0.41 |
|  | Males (n=94) | y= 0.20; x+5.3 | y= 0.54; x+5.3 | 0.79* | 0.62 |
| **Age** | Young (n=85) | y= 0.19; x+5.8 | y= 0.60; x+5.8 | 0.830* | 0.68 |
|  | Adult (n=130) | y= 0.19; x+5.3 | y= 0.60; x+5.3 | 0.751* | 0.56 |
|  | **Overall (n=215)** | **y= 0.18; x+5.4** | **y= 0.55 x+5.5** | **0.753\*** | **0.56** |

\*Significant correlation at $P \leq 0.01$ .
HbD= Hemoglobin determined spectrophotometrically; HbC= Hemoglobin calculated as 1/3rd of PCV; PCV= Packed cell volume

**Figure 1:**
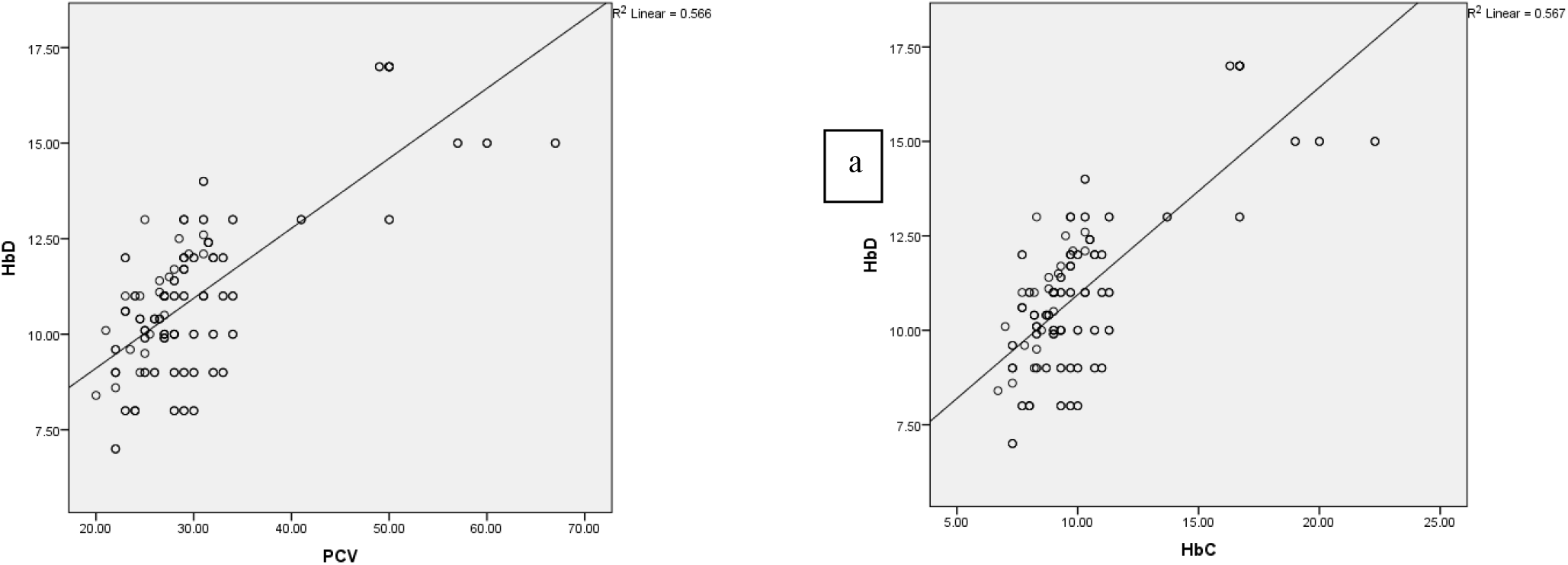
Scatterplot for logilinear regression between a) hemoglobin determined spectrophotometrically (HbD) and Packed Cell Volume (PCV) (n= 215; r= 0.753), and b) between hemoglobin determined spectrophotometrically (HbD) and hemoglobin calculated as one-third of Packed Cell Volume (HbC) (n= 215; r= 0.753) in Cholistani camels

## Discussion

In Pakistan, livestock is a sub-sector of agriculture and puts in about 56% to the agriculture value added services and almost 11% to the Gross Domestic Product (GDP). Pakistan has a large livestock population, well adapted to the local climatic conditions and has some of the best tropical dairy breeds. The livestock population in Pakistan comprises of about 53.8 million goats, 29.6 million cattle, 27.3 million buffalo, 26.5 million sheep and 0.9 million camels [17]. Despite of the fact that vegetation and the resources have been depleted, the livestock population is on the boom with the succession of years [12]. Cattle, sheep, goats and camels are the predominant types of livestock and the total population of livestock in Cholistan desert was estimated to be 12, 09, 528 heads, comprising of 47% cattle, 30% sheep, 22% goats and 1% camels [17]. In tropical pastoral system, in addition to shortage and poor quality of foodstuff, the decrease in the productive and reproductive potential of livestock can also be ascribed to the incidence of infections such as helminthiasis, trypanosomiasis, theileriosis, tick burden and tick borne infectivity. The parasitic infestation is one of the most important reasons of disease and production loss in the livestock by causing anemia and mostly death in heavily infected animals [18, 19, 20, 21]. Normally, for the diagnosis of anemia, PCV and Hb levels of the blood picture/complete blood count (cbc) are considered valid enough. In resource-poor settings (such as in Pakistan), automated veterinary hematology analyzers are not available. And human blood analyzers are usually being used for blood of livestock [22]. This poses the threat of erroneous results as the human analyzers are differently validated than the veterinary hematology analyzers [23]. This endorses the vitality of gold standard tests such as microhematocrit method (for PCV) and cyanmethemoglobin method (for Hb levels) [4, 15]. The present study included these two gold standard methods for deducing PCV and Hb in camel blood, and after attaining appropriate interrelationship between these attributes, puts forth a simple, pen-side hematological formula of Hb (g/dL)= 0.18(PCV)+5.4 for estimating Hb from PCV.

Considering the ‘hematological rule of three’ which is being implied in human medical practice, it has been well elucidated that Hb can be estimated as 1/3^rd^ of the PCV for apparently healthy human populations having normocytic normochromic erythrocytes [2, 24, 25]. On similar grounds, some studies on human blood have also negated the validity of this convention. In a malaria-endemic setting, this convention was not found valid in children and it was concluded that age, gender, season of sampling and physiological status of humans affects relationship between Hb concentration and PCV [7]. It was hence dubbed impossible to deduce a validated mathematical formula for their relationship as shown by other studies as well [6]. The earlier study dates back to 1994 which was conducted on human blood and endorsed that Hb was accurately measured as 1/3^rd^ of PCV and vice versa [26].

Regarding veterinary medical sciences, the only work reported on the interrelationship of PCV and Hb has been conducted on indigenous African cattle breeds and a formula of 0.28(PCV)+3.11 has been deduced for Hb estimation in g/dLs [9]. Similarly, pen-side hematological formulae for Hb estimation in goat blood [10] and for the blood of Cholistani breed of cattle have been reported by us earlier [11]. However, this is the first report on such interrelationship for camels being reared under pastoralism in Cholistan desert of Pakistan.

## Conclusions

Summing up, a convention of human clinical medicine that Hb concentration is a 1/3^rd^ of PCV and vice versa cannot be implied for the camels. However, a different equation *i.e*. Hb (g/dL) = 0.18(PCV)+5.4 may provide reliable results for Hb estimation from the PCV in this specie. The results of the study may be of substantial value to the researchers, academicians and veterinary clinicians of resource-poor areas. It is suggested that other mathematical formulae regarding hematological attributes being used in human clinical medicine may also be validated for various use in veterinary medical practice.

**Figure 2:**
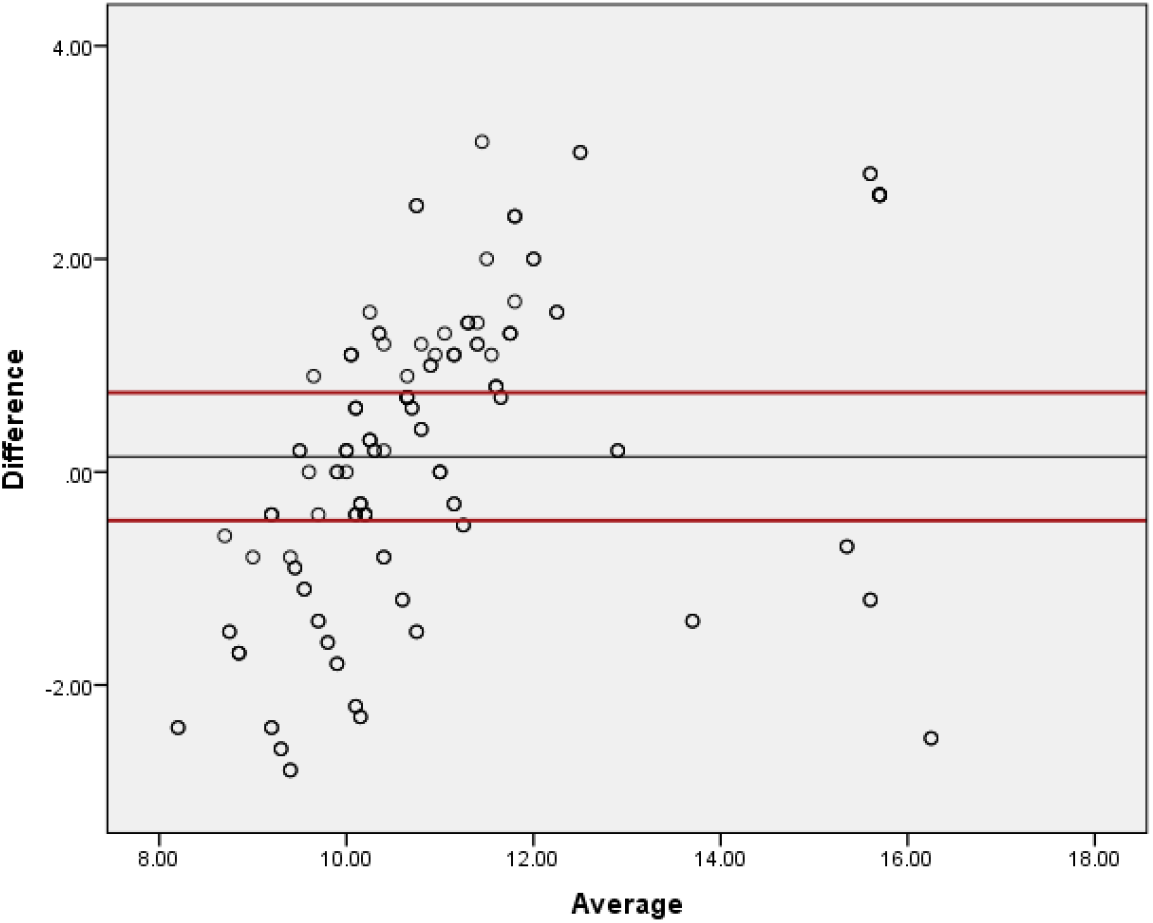
Scatterplot of Bland and Altman test between difference of hemoglobin determined spectrophotometrically and corrected hemoglobin (HbD-CHbC) and average of both hemoglobins (HbD+CHb/2) in Cholistani camels (n= 215) Black line indicates mean difference (0.1436) whereas the upper and lower red lines indicate upper (3.00) and lower (-2.72) values for 95% CI

## Acknowledgements

The authors are grateful to the ‘**Pakistan Science Foundation**’ for provision of research grant No. PSF/NSLP/P-IUB-931 titled “Devising and Validating Pen-side Hematological Tests as an Enhanced Approach to the Diagnosis of Anemia in Cholistani Livestock (Camels and Cattle)”.

## Author Contributions

**Conceptualization**: Umer Farooq and Mushtaq Lashari

**Data curation:** Shahbaz Ahmad and Haroon Rashid

**Formal analysis:** Nouman Sajjad, Musadiq Idris and Aisha Mahmood

**Investigation:** Nouman Sajjad

**Methodology:** Shahbaz Ahmad, Musadiq Idris, Zia Rehman and Aisha Mahmood

**Project administration:** Umer Farooq

**Resources:** Mushtaq Lashari

**Software:** Zia Rehman and Haroon Rashid

**Validation:** Musadiq Idris

**Writing:** Sajid Hameed;

**Review and editing:** Sajid Hameed

